# Indirect effects of habitat amount mediated by habitat configuration determine bat diversity at the landscape-scale in Peninsular Malaysia

**DOI:** 10.1101/2023.12.30.573737

**Authors:** Quentin C.K. Hazard, Natalie Yoh, Jonathan Moore, Juliana Senawi, Luke Gibson, Jérémy S.P. Froidevaux, Ana Filipa Palmeirim

**Affiliations:** CIBIO, Centro de Investigação em Biodiversidade e Recursos Genéticos, InBIO Laboratório Associado, Campus de Vairão, Universidade do Porto, 4485-661 Vairão, Portugal; BIOPOLIS Program in Genomics, Biodiversity and Land Planning, CIBIO, Campus de Vairão, 4485-661 Vairão, Portugal; Durrell Institute of Conservation and Ecology, University of Kent, Canterbury, CT2 7NR, UK; School of Environmental Science and Engineering, Southern University of Science and Technology, Shenzhen, China; School of Environmental Sciences, University of East Anglia, Norwich, UK; Department of Biological Sciences and Biotechnology, Faculty of Science and Technology, Universiti Kebangsaan Malaysia, 43600 Bangi, Selangor, Malaysia; Langkawi Research Centre, Campus of Tunku Abdul Halim Mu’adzam Shah, Universiti Kebangsaan Malaysia, Jalan Teluk Yu, Kampung Kok, 07000 Langkawi, Kedah, Malaysia; University of Stirling, Biological and Environmental Sciences, Faculty of Natural Sciences, Stirling, UK.; Centre d’Ecologie et des Sciences de la Conservation (CESCO, UMR 7204), CNRS, MNHN, Sorbonne-Université, Concarneau/Paris, France

**Keywords:** Chiroptera, Habitat loss, Fragmentation, Structural Equation Modelling, Passive acoustic monitoring, Habitat Amount Hypothesis

## Abstract

The impacts of fragmentation are often overlooked in landscape studies investigating how habitat loss impacts biodiversity, despite the casual relationship linking both processes. As habitat loss is the primary cause of fragmentation, understanding the inter-related effects of these twin processes on biodiversity is key to minimise biodiversity loss. Here we assess how habitat amount and configuration influence insectivorous bat assemblages, considering both the direct effects of these processes, as well as the indirect effects of habitat amount mediated through configuration. Bats were acoustically surveyed along independent gradients of habitat amount (forest cover) and configuration (number of patches and edge density) across 28 insular landscapes embedded within a Malaysian hydroelectric reservoir. Using Structural Equation Modelling, the direct and indirect effects of habitat amount were examined on bat sonotype richness, total, and guild-specific activity (forest, edge and open-space foragers). Forest cover had a direct and positive effect on sonotype richness and forest forager activity. The quadratic relationship linking edge density and forest cover was strong and overall positive, but while below 30% of forest cover, increasingly forested landscapes had increasingly high edge densities, the opposite pattern was observed in more forested landscapes. On the other hand, increasingly forested landscapes consistently harboured less patches. Owing to the overall low habitat amount in our landscapes and negative edge effects, the indirect effects of forest cover (mediated through edge density) were therefore negative on sonotype richness, outweighing any positive direct effect. The number of patches had little effect on the bat assemblage, except on total activity which was higher in landscapes harbouring more forest patches. As a result, negative indirect effects of forest cover mediated through number of patches were only observed on total activity. Our results highlight that, in natural settings, habitat amount can hardly be altered without influencing habitat configuration, thereby preventing any independent management of these threats. Minimising habitat loss is therefore essential to balance the associated prevailing negative effects of fragmentation on insectivorous bats across tropical forests.

## 1. Introduction

The detrimental effects of habitat loss – a reduction in the habitat amount – on biodiversity have been widely acknowledged in the literature (Arasa-Gisbert et al. 2022; Caro et al. 2022; Sánchez-Bayo & Wyckhuys 2019). However, the effects of fragmentation are still widely debated (Miller-Rushing et al. 2019). Indeed, while many studies conducted at the local scale rely on the effects of patch size and isolation to infer negative effects of fragmentation (i.e., the change in habitat configuration as a continuous habitat is split into several smaller pieces) (Haddad et al. 2015), Fahrig (2013) suggests that by failing to control for the total amount of habitat in the landscape, such studies do not disentangle the effects of fragmentation from those of habitat loss. Fahrig (2003, 2017) further suggests that when considered independently from habitat loss, fragmentation “per se” (i.e, the change in habitat configuration as a continuous habitat is split into several smaller pieces *while keeping the total habitat amount in the landscape constant*) often has unimportant or even positive effects on species richness.

In natural settings, habitat loss is the primary source of fragmentation (Cushman et al. 2012; Hanski 2015; Liu et al. 2016). As any change in habitat amount tends to alter habitat configuration, either by influencing the number of patches or the edge density (Didham et al. 2012), strong collinearity often exists between habitat amount and the most commonly used configuration metrics (e.g., number of patches, mean patch size, mean inter-patch distance and edge density (Wang et al. 2014)). Yet, classic statistical methods such as multiple regressions are not tailored to handle such collinearity, often leading to significant biases when investigating the relative importance of both predictors in explaining biodiversity patterns (Ruffell et al. 2016; Smith et al. 2009). However, by accounting for the effects of habitat amount and habitat configuration on a given response, along with those of the former on the latter, Structural Equation Modelling enables us to quantify their respective direct effects, as well as the indirect effects of habitat amount mediated through configuration therefore accounting for any collinearity between both predictors (Didham et al. 2012; Ruffell et al. 2016). This method further allows the reconciliation of formerly conflicting perspectives by simultaneously accounting for and quantifying the effects of habitat loss (being considered here as a reduction in the habitat amount independent from habitat configuration), fragmentation *per se* (direct effects of habitat configuration, independent from habitat amount), and fragmentation (considered here as the indirect effects of habitat amount mediated through configuration) (Figure 1).

**Fig. 1.**
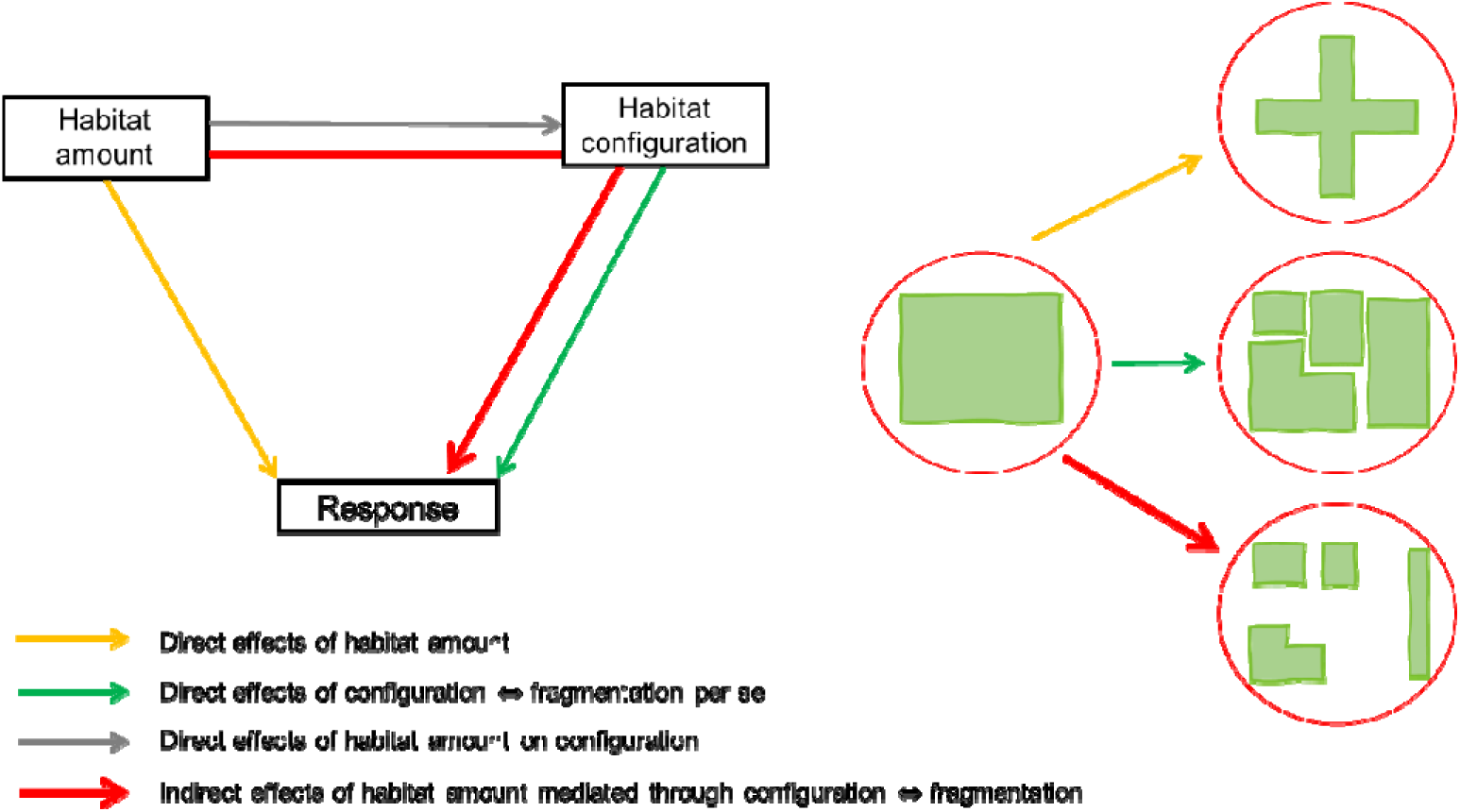
Representation of the interplay between habitat amount and configuration. Structural Equation Modelling can be used to assess the effects of three potential drivers of biodiversity response: (i) habitat loss (a reduction in the habitat amount independent from habitat configuration, i.e. without affecting edge density or number of patches), (ii) fragmentation *per se* (a change of habitat configuration independently from habitat amount), and (iii) fragmentation (a change in the habitat amount further affecting the habitat configuration).

Furthermore, whilst strong, the relationship between habitat amount and configuration might be non-linear in natural settings (Püttker et al., 2020). In this case, below a certain threshold of habitat loss, removing habitat tends to increase the number of patches and edge density in the landscape. Conversely, beyond this threshold, habitat removal is instead generally associated with a decrease in edge density and the number of patches (Figure 2) (Villard & Metzger 2014). This complex relationship challenges the long-held view that habitat loss and fragmentation are independent processes that can be managed independently (Smith et al. 2011). However, there are few studies shedding light on the intertwined effects of habitat amount and configuration (but see for instance Cosentino & Brubaker (2018), Püttker et al. (2020), and Suárez-Castro et al. (2020)).

**Fig. 2.**
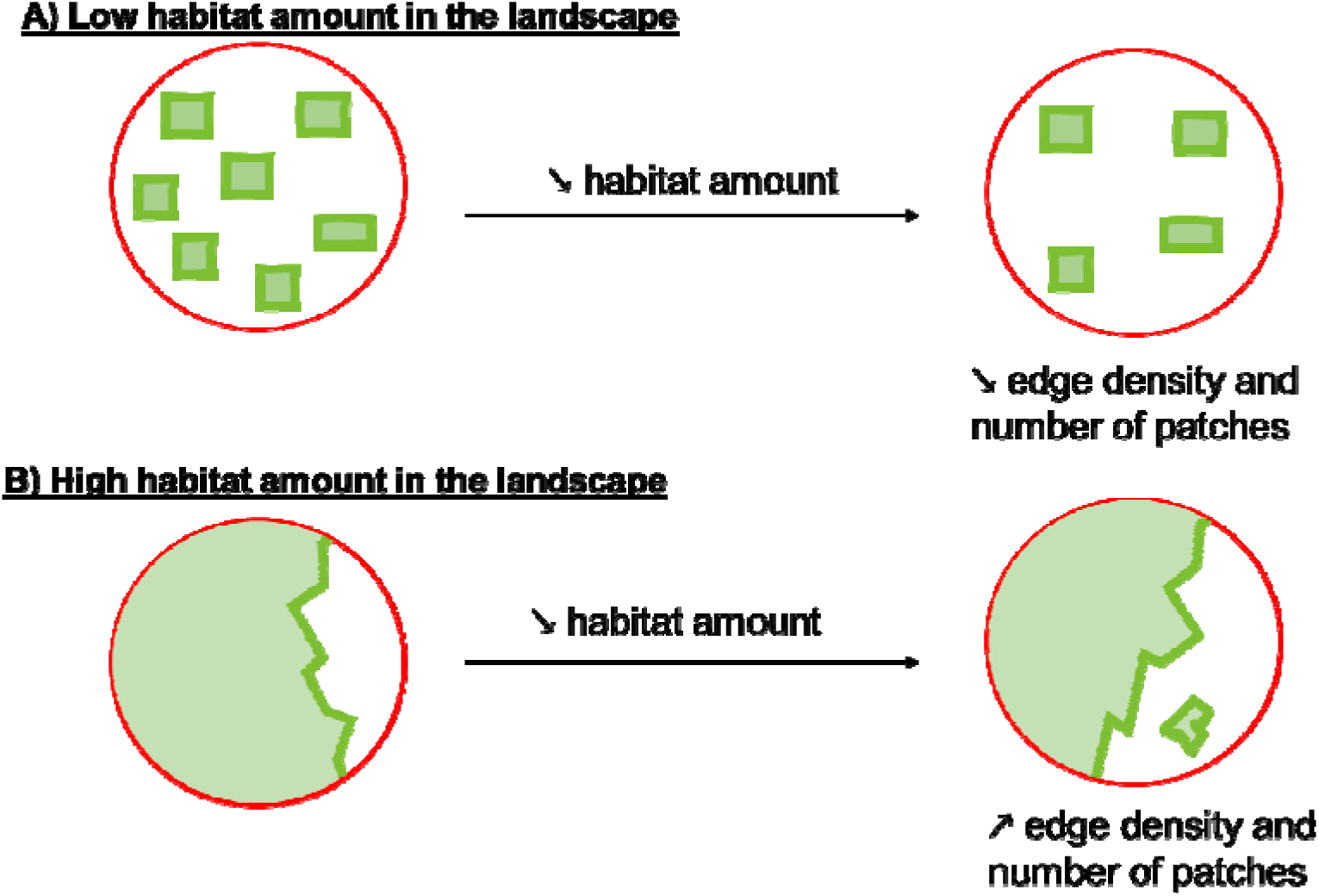
Expected scenarios in natural settings for changes in the fragmentation *per se* following a decrease in the habitat amount of landscapes harbouring A) a low (<20%) or B) high habitat amount (>50%). In A), a decrease in the habitat amount is more likely to result in a decrease in both the number of patches and edge density, except for some specific cases (Villard & Metzger 2014). In B), a decrease in habitat amount is more likely to increase the number of patches and edge density. Circular landscapes are delimited in red, within which habitat is shown in light green and edges in dark green.

The effects of habitat loss and fragmentation might further vary according to species-specific traits such as body size, trophic level, or mobility (Ewers & Didham 2006). For instance, the idiosyncratic responses shown by different bat species to the twin processes of habitat loss and fragmentation are a function of dietary specialisations (Meyer & Kalko 2008), wing morphologies (Bader et al. 2015; Farneda et al. 2015; Furey & Racey 2016), roosting habits (Struebig et al. 2008, 2009), and echolocation calls structures (Hazard et al. 2023; López-Bosch et al. 2021). Given their detection function, echolocation calls are modulated by a bats’ habitat and foraging preferences (Denzinger & Schnitzler 2013; Schnitzler & Kalko 2001). Facing antagonistic physical constraints, forest, edge, and open-space foragers have evolved calls designed for prey detection in these specific environments, and their distinctive call signatures allow for the differentiation of these three foraging guilds (Yoh, Kingston, et al. 2022). Specifically, when foraging, forest bats face the challenge that prey-returned echoes are masked by those returned by vegetation clutter (Arlettaz et al. 2002). Forest-foraging bats therefore emit precise detection calls over short distances. On the other hand, open-space foragers have to locate prey in open spaces, emitting less accurate but long-range echolocation calls, while edge foraging bats emit calls that enable them to locate prey against background features at intermediate distance (Denzinger & Schnitzler 2013; Schnitzler & Kalko 2001). Forest bats are therefore expected to persist and be more active in more forested landscapes, while open-space or edge foragers may favour sparsely forested or edge-dominated landscapes.

In this study, we aimed to disentangle the direct effects of habitat amount from its indirect configuration-mediated effects on Malaysian insectivorous bat assemblages across a hydroelectric reservoir. By flooding lowland areas, damming creates a myriad of insular forest fragments of various sizes and isolations, sitting on a uniformly inhospitable water matrix. Hydroelectric reservoirs therefore represent unique experimental settings to study the effects of habitat amount and configuration without confounding matrix or strong local-history effects (Diamond 2001; Palmeirim et al. 2022; Semper-Pascual et al. 2021). Using acoustic detectors, we surveyed bats across 28 landscapes of varying forest cover, edge density and number of patches at the landscape-scale. As bats feeding under similar environmental constraints often emit similar calls, we classified calls into sonotypes, i.e., mixed-species groups of similarly shaped calls (Yoh 2018), rather than to the species level. Each sonotype was further classified according to its habitat affinity (i.e. forest, edge or open-space forager). We hypothesised bat diversity to be directly and positively affected by habitat amount. Given the strong evidence of the pervasiveness of edge effects, especially in high contrast matrix types (Ewers & Didham 2006; Pfeifer et al. 2017), we anticipated edge density to negatively impact richness and total activity. Conversely, by promoting habitat heterogeneity, the presence of a multitude of patches may also allow disturbance-tolerant species to occupy a niche hitherto unavailable to them (Brändel et al. 2020; Palmeirim et al. 2019): the number of patches was therefore expected to positively influence sonotype richness. We also hypothesised that the response of bats would be mediated by their foraging guild, with fragmentation *per se* benefitting edge and open-space foragers while disfavouring forest-dependent bats, the latter also being expected to be negatively impacted by habitat loss (Hazard et al. 2023; López-Bosch et al. 2021; Rocha et al. 2018). Considering the high deforestation level in most of the studied landscapes (<30% of forest cover), we anticipated more forested landscapes to harbour a higher edge density and number of patches compared to less forested landscapes, therefore resulting in the indirect effects of habitat amount mediated through configuration being positive on richness, total activity, and on the activity of edge and open-space foragers but negative on that of forest foragers.

## 2. Material and methods

### 2.1 Study area

Surveys were carried out at Kenyir Lake in Northeast Peninsular Malaysia (Figure 3). This 37 years old freshwater hydroelectric reservoir was formed by the damming of the Kenyir River, resulting in the flooding of nearly 36,900 ha of forest, fragmenting the region into over 340 insular patches. These former hilltops of lowland and mid-elevation dipterocarp forest used to be selectively logged prior to the creation of the dam (Muhammad Yusuf 2005; Qie et al. 2011; Yong 2015). Annual precipitation ranges between 2700 and 4000 mm, and this region undergoes a rainy season from November to March, and a dry season from May to October (Qie et al. 2011).

**Fig. 3.**
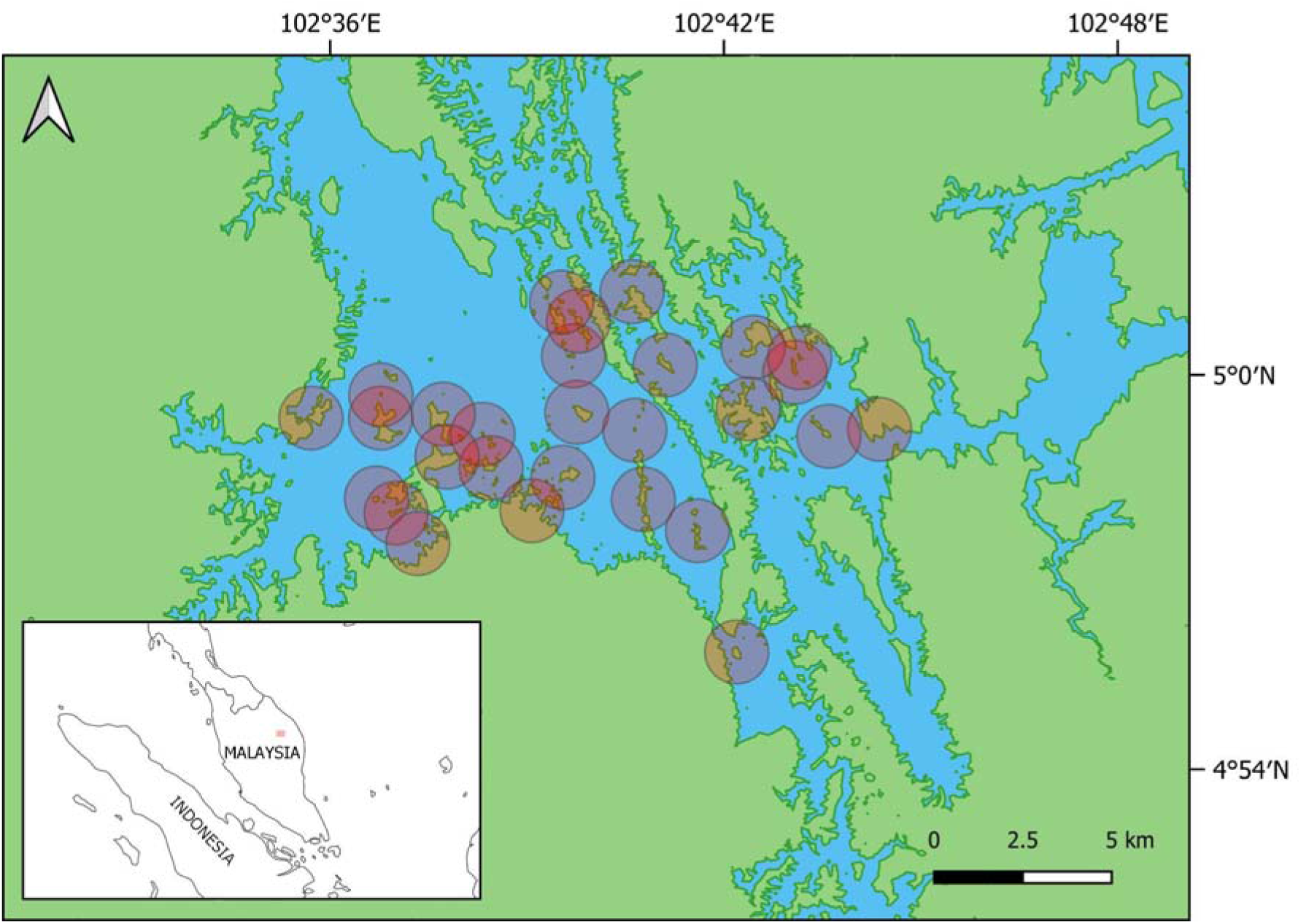
Location of the study area and sampled landscapes in the Kenyir Lake, Malaysia. The 900 m-radius landscapes – the largest landscape size considered in this study – centred on the sampling site are displayed in pink. The size of the different buffers used throughout the analysis are available in Table 1. Mainland continuous forest and insular forest patches are shown in *green*, and the aquatic matrix in *blue*.

**Table 1.**
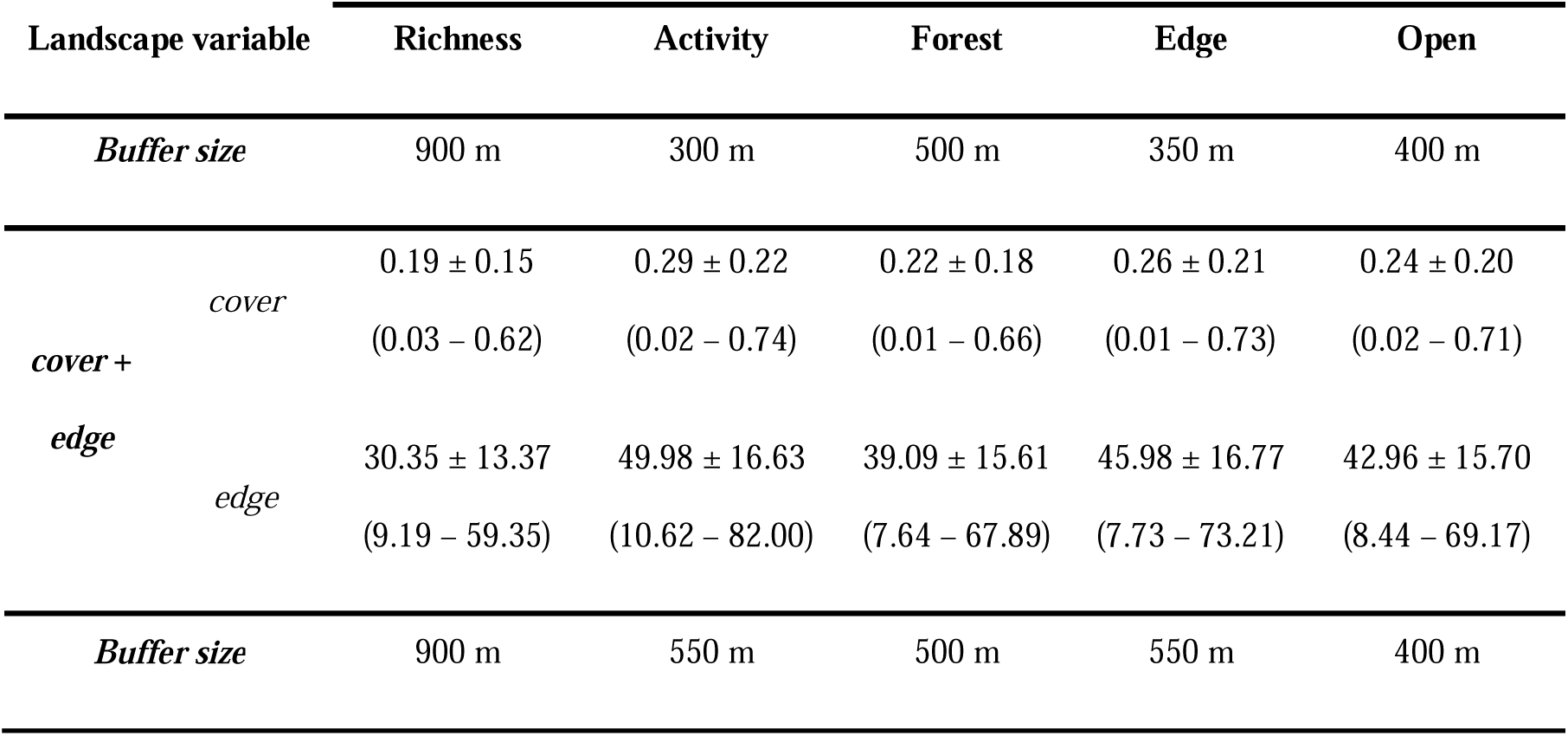
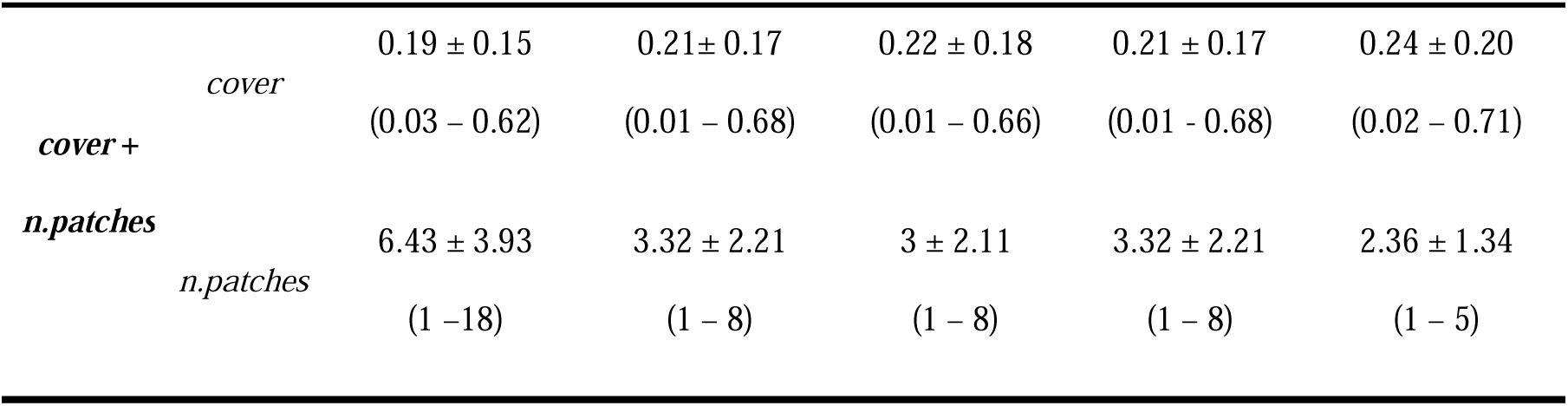
Buffer sizes selected based on the analysis of the scale of effect of both habitat amount (*cover*) and configuration (*edge* or *n.patches*) for each bat diversity response – sonotype richness, total activity and activity of forest, edge and open-space foragers. The scale of effect was analysed considering both combinations: (1) *cover* and *n.patches* and (2) *cover* and *edge*. The AICc of the models for each considered scale can be found in Figure S2. For the “best” scale of each response variable, we show the mean (min – max) of the explanatory variables at the given scale. Variation in each of the explanatory variables for the remaining considered scales considered is shown in Figure S1.

### 2.2 Data collection

We selected 28 sampling sites buffered by heterogeneous levels of forest amounts and configurations. Insectivorous bats were acoustically surveyed between September 8^th^ and October 13^th^ 2019, a period during which heavy rainfall is infrequent, thereby ensuring the quality of the acoustic recordings. We deployed one Audiomoth© recorder (Hill et al. 2018) per sampling site and simultaneously sampled between one and eight sites, depending on fieldwork constraints (e.g. site accessibility). At each sampling site, recorders were set at a sampling rate of 384 kHz and medium gain and placed in the forest interior between 14 and 123 m (median: 50 m) from the forest edge, at 2 m above ground. The recordings were made on a single occasion. As to cover bat activity peaks, recordings were taken at dawn and dusk for six hours: first from 6 pm to 10 pm (i.e., starting 30 minutes before sunset), and the second from 4 am to 6 am (i.e., ending 30 minutes after sunrise) (Mariton et al. 2023). A single visit to each site does not allow to fully capture the local bat assemblages, but we adopted this approach to maximise the number of sites surveyed while keeping the acoustic data management process low (López-Baucells et al. 2021). As the aim of the study was to assess the response of bat assemblages to habitat modification, the sampling effort implemented proved to be sufficient to detect an effect (Hazard et al. 2023; López-Bosch et al. 2021).

### 2.3 Acoustic analysis

Files containing sounds ranging between 10 and 250 kHz, with a minimum pulse length of 2 ms and a maximum pulse length of 500 ms were filtered (Altringham 2011) using Kaleidoscope software (Wildlife Acoustics 2019). A sound event was counted as a bat pass if two or more pulses of a single sonotype were detected on a five-second acoustic file (Yoh et al. 2023). Bat calls were further automatically classified into one of the eight sonotypes: Frequency Modulated (FM), Frequency Modulated quasi–Constant Frequency (FMqCF1 to FMqCF5), Quasi Constant Frequency (QCF), Constant Frequency (CF), as supported by the classifier developed by Yoh et al. (2022). This automatic classification was followed with a manual verification (López-Baucells et al. 2019). CF sonotypes could further be identified to the species level based on the reference calls compiled in Hazard et al. (2023). The FMqCF1 sonotype described in Yoh et al. (2022) was not detected in our dataset. LF, FMqCF2 and FMqCF3 calls were classified as belonging to open-space foragers, FMqCF4, FMqCF5 and QCF to edge foragers, and FM and CF calls to forest foragers (Yoh, Kingston, et al. 2022) (see Hazard et al. (2023) for further information about the sonotype classification).

### 2.4 Landscape variables

Percentage of forest cover [*cover*] indicated habitat amount, and both forest edge density (i.e., ratio between the total forest/water interface in the landscape and the total landscape area, m/ha) [*edge*] and number of forest patches [*n.patches*] indicated habitat configuration. Proximity of the sampling site to forest edge was accounted for using the Euclidean distance between the detector and the nearest forest edge *[near.dist]*. These metrics were calculated utilising the “landscapemetrics” package (Hesselbarth et al. 2019) (see Hazard et al. (2023) for further details on the calculation of these metrics). Landscape variables were calculated for 16 circular buffer sizes ranging from 250- to 1000-m radii, with 50-m increments, and whose centroids were matching the location of each acoustic detector (Hesselbarth et al. 2019) (Figure S1). This size scope aimed to consider the heterogeneity in bat home ranges (Jackson & Fahrig 2012), further including a range of habitat amounts and configurations, while minimising the overlap between contiguous buffers (Zuckerberg et al. 2020). For landscapes smaller than 400-m radii, the distribution of *n.patches* did not include much variation (1 to 4 patches) (Figure S1). As such, this variable was only considered in the subsequent analyses for buffers larger than 400-m radii.

### 2.5 Data analysis

#### 2.5.1 Response variables

We considered five response variables to evaluate bat responses to habitat amount and configuration, namely sonotype richness, total bat activity, and guild-specific activity (i.e., forest, edge, and open-space foragers). We used sonotype richness, i.e. the number of sonotypes, as a proxy for bat species richness, and bat activity (i.e., number of bat passes per night) as a surrogate of bat abundance (Hazard et al. 2023; López-Bosch et al. 2021). Both total and guild-specific activities were consistently log-transformed through the analyses, to meet normality assumptions. All models were run using a gaussian error distribution: negative binomial distribution was also considered but was outperformed by the Gaussian error structure when a log transformation was applied.

Because they could not be matched with their corresponding sonotypes, social calls were accounted for in the total activity but were neither identified to sonotype nor guild levels. Potential spatial autocorrelation was examined using a Mantel test that compared geographic distances and differences in sonotype richness, total and guild-specific activity between all possible pairwise site combinations. The absence of spatial autocorrelation was systematically confirmed when testing such correlation on model residuals (Mantel test, *p* < 0.05). Pairwise geographic distances were calculated using the R package “geodist” (Karney 2013), and the Mantel test was performed using the R package “ade4 (Dray & Dufour 2007).

#### 2.5.2 Scale of effect

We examined the best “scale of effect”, i.e., the buffer size that most strongly affects the focus response (*sensu* Jackson & Fahrig (2012)). To do so, we fitted Linear Models (GLMs) with a Gaussian error distribution between each of the five response variables and each pair of habitat amount (*cover*) and fragmentation variables (*n*.*patches* or *edge*) for each of the 16 buffer sizes. This allowed us to determine at what scale the two variables had the strongest joint effect on each of the response variables, i.e., the lowest Akaike Information Criterion corrected from small samples sizes (AICc) to both landscape variables considered among all the fitted models (Jackson & Fahrig 2012; Püttker et al. 2020; Rios et al. 2021) (Figure S2). When examining the scale of effect for the activity of forest foragers as influenced by *cover* and *n.patches*, the models regarding the buffers of 350-, 400- and 950-m radii did not converge: these buffer sizes were therefore not considered in the scale selection for this response variable.

#### 2.5.3 Interrelated effects of habitat loss and fragmentation

We then examined the interrelated effects of habitat loss and fragmentation on bat assemblages using Structural Equation Models (SEMs). SEMs are to date one of the most adequate statistical tools when addressing the relative effects of these processes, accounting for relationships among causally linked collinear variables (Didham et al. 2012; Ruffell et al. 2016). To separate the direct from the indirect effects of habitat loss, we designed piecewise SEMs based on the assumptions that 1) the response variables would be influenced directly by both fragmentation and habitat amount, and that 2) the causal relationship between habitat amount and configuration would cause habitat amount to indirectly affect the response through its effects on habitat configuration (Bollen & Pearl 2013; Pearl 2012; Püttker et al. 2020). Only one configuration metric was used at a time: each response variable was therefore analysed twice, first considering *edge* and then *n.patches*. In order to account for the possible influence of the distance at which the acoustic detector was deployed from the edge, [*near.dist*] was added in each model as a design covariate. While the relationship between *cover* and *n.patches* in a given set of same-sized landscapes was consistently linear, the relationship between *edge* and *cover* was bell-shaped. In this case, as integrating the quadratic term of *cover* consistently decreased the full model’s AICc, it was added in all models including *edge* as the fragmentation metric (Fahrig 2003; Villard & Metzger 2014). The basis set, i.e. models making up the SEM, was therefore compounded of two coupled linear models, the first relating the response with *cover*, configuration (either *edge*, or *n.patches*), and *near.dist*, and the second relating configuration with *cover* (and *cover²*, in the case of *edge* being the configuration variable). In each basis set, both cover and each of the configuration variables were included at their best joint scale of effect (Figure S2). A gaussian error structure was used to fit the models. Prior to running the models, total and guild-specific activity were log-transformed to meet normality assumptions, and all the variables were standardised to enable the comparison of effect size. The piecewise SEM were carried out using the R package “piecewiseSEM” (Lefcheck 2016). We further computed the piecewise SEM models in the semEff function from the “semEff” R package (Murphy 2022) to calculate the direct and indirect effects of the predictors on the responses, along with their parametric, accelerated bias-corrected bootstrapped confidence intervals to assess their statistical significance (Palmeirim et al., under review). Indirect effects of the percentage of forest cover through configuration were calculated by multiplying the path coefficient of the effect of the percentage of forest cover on configuration by the path coefficient of the effect of configuration on richness (Grace 2007). The bootstrapped confidence intervals were generated using credible intervals of 0.95 with 9999 iterations (Murphy 2022).

We confirmed the assumptions regarding the distribution of the variables and their residuals by running each model into the “performance” (Lüdecke et al. 2021) and “DHARMa” (Hartig 2022) R-packages. Inter-variable correlation was low to moderate (|*r*| < 0.70). A basis set was examined only if Fisher’s C test of directed separation was not significant. This test examines the different components of the network and determines whether the non-tested paths are important in explaining the data, therefore depicting the overall relevance of specified pathways (Lefcheck 2016; Shipley 2000). All data analyses were performed in R (R Core Team 2022).

## 3. Results

### 3.1. Bat activity

We obtained a total of 47,324 five-second recordings, with almost half of them (21,197) containing bat passes. From these, 16 sonotypes were identified, corresponding to nine sonotypes matching a single species and seven sonotypes grouping multiple species. Ten sonotypes were classified as forest foragers, three as edge foragers, and three as open-space foragers (Table S1). Sonotype richness and activity were highly variable between sites, ranging from four to 13 sonotypes, and from 43 to 3,351 bat passes recorded. Edge foraging bats showed the highest activity (69.3%), in comparison to open-space (18.5%) and forest foragers (14.4%). Indeed, the most often recorded sonotype was the edge forager FM4, accounting for 66.8% of the bat passes. In total, 989 passes, mostly social calls (99.2%), were not identified to the guild level, and were only included in the total activity.

### 3.2. Scale of effect

The scale of effect varied according to the response variable considered (Table 1). Sonotype richness presented lower AICc values at a broader scale (900 m when considering *cover* with either *edge* or *n.patches*), whereas AICc values for total activity, forest, edge, and open-space foragers activity were lower for smaller scales (300, 500, 350 and 400 m, respectively, when considering *cover* and *edge*, and 550, 500, 550 and 400 m respectively when considering *cover* and *n. patches*). Interestingly, the scale of effect of the different response variables were relatively close among the two fragmentation variables (Table 1).

### 3.3. Influence of habitat configuration on the habitat amount

Habitat amount was generally low across the landscapes, the average *cover* being consistently <30% (Table 1). The non-linear relationship linking *cover* and *edge* was, overall, positive (Figure 4A, Table S2). *Cover* was a good predictor for *edge*, with *R*² ranging between 0.56 and 0.61.

**Fig. 4.**
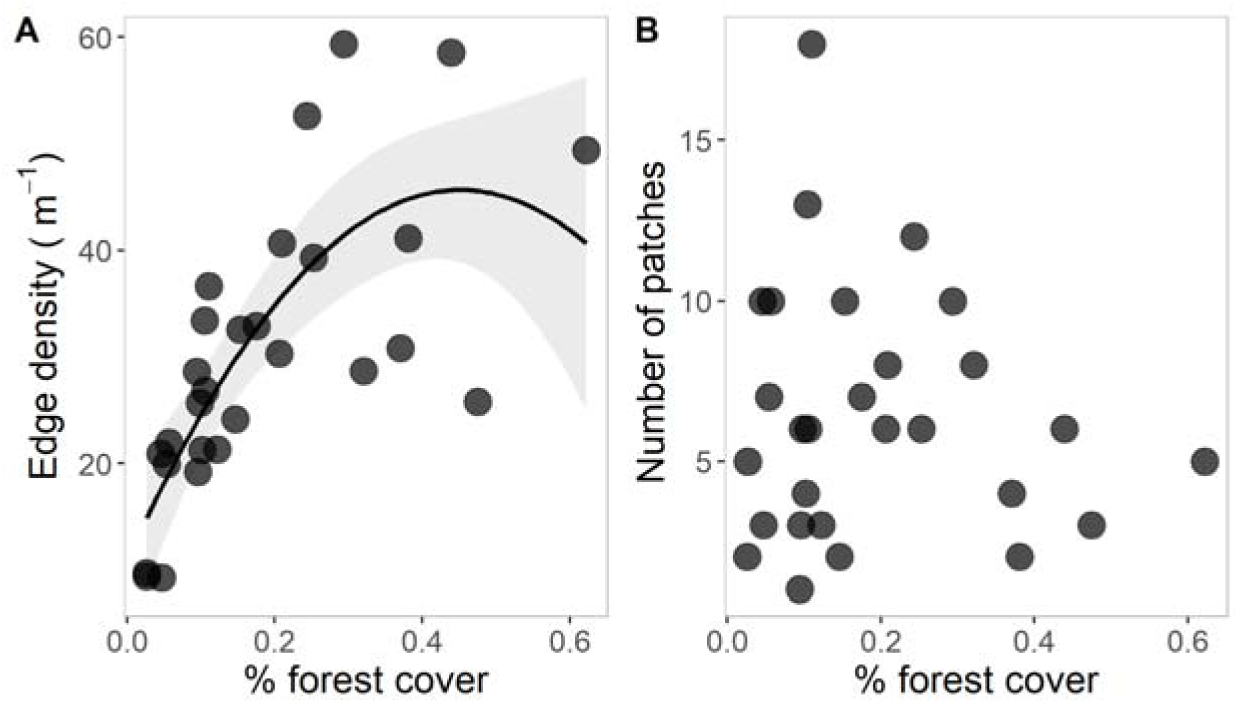
Relationship between the percentage of forest cover and A) edge density, and B) number of patches across the 28 landscapes with 900-m radius surveyed in the Kenyir Lake. The solid line depicts the prediction given by the linear model for significant relationships, and the shaded area shows the 95% confidence interval.

The relationship between *cover* and *n.patches* was negative in all the considered models, except for the one with a 900-m radii landscape where it was non-significant (Figure S3B, Table S3).

### 3.4 Direct and indirect effects of forest cover as mediated by edge density

When using *edge* as the habitat configuration variable, forest cover had a positive direct effect on sonotype richness (*β* = 0.476 ± 0.146, CI_min_ = 0.200, CI_max_ = 0.735). Edge density had a negative effect on sonotype richness (*β* = –0.422 ± 0.149, CI_min_ = –0.693, CI_max_ = –0.110), total bat activity (*β* = 0.553 ± 0.126, CI_min_ = – 0.741, CI_max_ = –0.263) and edge sonotype activity (*β* = –0.520 ± 0.124, CI_min_ = –0.740, CI_max_ = –0.288). Given the positive relationship between *cover* and *edge* (Figure 4), *cover* also indirectly and negatively influenced sonotype richness (*β* = –0.218 ± 0.152, CI_min_ = –0.451, CI_max_ = −0.053), total bat activity (*β* = –0.392 ± 0.108, CI_min_ = –0.587, CI_max_ = –0.182) and edge sonotype activity (*β* = –0.343 ± 0.103, CI_min_ = –0.590, CI_max_ = –0.182) (Figure 5).

**Fig. 5.**
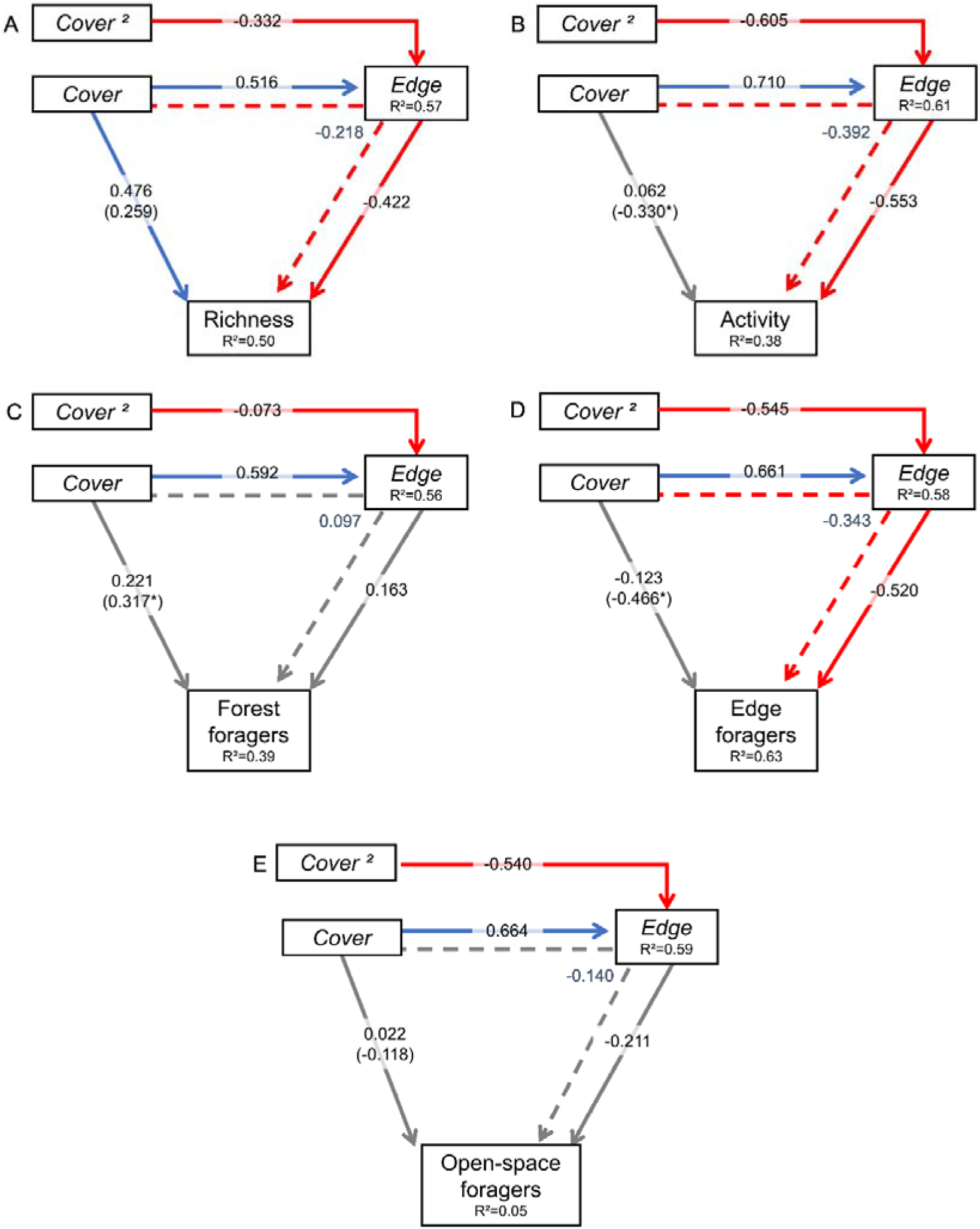
Results of the Piecewise Structural Equation Models representing the effects of habitat amount (*cover* and *cover²*) and edge density on sonotype richness, (b) total activity (log_10_ *x*), and activity of (c) forest sonotypes (log_10_ *x*), (d) edge sonotypes (log_10_ *x*), and (e) open-space sonotypes (log_10_ *x*). Blue arrows depict positive relationships, red arrows depict negative relationships, and grey arrows depict non-significant relationships.

Solid arrows depict direct relationships, and dashed arrows represent indirect relationships. For each response variable, we indicated the standardised path coefficients. The Euclidean distance point-edge was omitted for clarity as it was never significant. Coefficients for this variable can be found in Table S2.

### 3.5. Direct and indirect effects of forest cover as mediated by the number of patches

When using *n.patches* as the habitat fragmentation variable, forest cover had a positive direct effect on the activity of forest sonotypes (*β* = 0.399 ± 0.148, CI_min_ = 0.062, CI_max_ = 0.647). The number of patches positively affected total bat activity (*β* = 0.428 ± 0.078, CI_min_ = 0.146, CI_max_ = 0.692). The pathway analysis explained a large amount of variation, *R*² ranging between 0.23 for open-space sonotypes and 0.44 for the activity of forest foragers. Given the negative relationship between forest cover and the number of patches, forest cover had a negative indirect effect on total bat activity (*β* = –0.161 ± 0.078, CI_min_ = –0.349, CI_max_ = –0.033) (Figure S3). Here, the amount of variation explained varied substantially among the response variables being considered, with the coefficient of determination ranging between *R*² = 0.07 for open-space foragers, and *R*² = 0.44 for forest foragers.

## 4. Discussion

While an extensive body of literature has aimed to unravel the relative importance of habitat loss and fragmentation in driving changes in species diversity over the past decades, most landscape-scale studies do not control for the causal relationship linking both processes (Didham et al. 2012; Ruffell et al. 2016). Here we helped fill this gap by examining the direct effects of habitat amount and indirect effects as mediated by configuration on Southeast Asian insectivorous bat assemblages across highly deforested landscapes. Overall, we found that the effects of fragmentation *per se* were mostly negative, but dependent on the metric used (i.e., edge density or number of patches). The influence of habitat amount manifested both directly and indirectly via its impact on habitat configuration. Although the direct effects of habitat amount on the bat assemblage were overall positive, habitat amount was positively associated with edge density and negatively with the number of patches. As a result, the indirect effects of habitat amount mediated through edge density were negative on bat sonotype richness and activity, but positive on bat activity when mediated through number of patches. Our results suggest that habitat amount and configuration are two interrelated sides of a global process, with habitat amount acting on the response both directly and indirectly by influencing habitat configuration (Villard & Metzger 2014).

### 4.1 Direct effects of habitat configuration

Our results do not support the hypothesis that the effects of fragmentation *per se* are mostly unimportant, or positive whenever significant (Fahrig 2003, 2017). We found the effects of fragmentation *per se* to be mostly negative, but dependent on the fragmentation metric used: edge density negatively influenced sonotype richness, total activity, and edge sonotype activity, while the number of patches positively influenced total bat activity.

While forest patches usually harbour a lower bat richness than continuous forest in the Neotropics (Cosson et al. 1999; Rocha et al. 2017; Schulze et al. 2000) and in the Paleotropics (Struebig et al. 2008), landscape-scale studies have highlighted that bat abundance (Ethier & Fahrig 2011) and richness (Farneda et al. 2020; Meyer & Kalko 2008) tend to benefit from fragmentation. Indeed, in low-contrast landscapes such as primary/secondary vegetation matrixes, forest edges represent linear delimitations between two habitats and generally harbour a lower structural complexity compared to surrounding forest-interiors, often supporting high arthropod abundance: these habitats are therefore well suited for insects-feeding bats (Verboom & Huitema 1997). Consistently, in temperate regions (Ethier & Fahrig 2011; Müller et al. 2012; Wolcott & Vulinec 2012) and in the Neotropics (Delaval & Charles-Dominique 2006), bat activity has been reported to be higher along forest edges compared to surrounding matrix habitats. Yet, in the context of insular fragmented landscapes, edge density depicts the delineation between terrestrial habitat (here forest) and freshwater habitat (here open water). With the exception of a few species (e.g., *Miniopterus magnater, Myotis hasseltii*, *Cynopterus horsfieldii*, see Lim et al. (2017)) bats are not known to forage above open water in our study area; as such, the negative effects of edge density on total bat activity are likely the result of this high forest-matrix contrast. Moreover, even edge-adapted bats showed a negative response to edge density, which was unexpected considering that they have been reported to be unaffected or positively affected by edge effects (Froidevaux et al. 2022; Yoh, Clarke, et al. 2022), and tolerant to forest disturbances, such as logging (Struebig et al. 2013). In addition to their edge-adapted morphology (high aspect ratio and low wing loading) (Norberg & Rayner 1987), the calls emitted by edge foragers are mostly tailored for navigation and prey detection in partially cluttered habitats such as forest edges, where they therefore tend to be more abundant (Meyer et al. 2004). In the case of an aquatic matrix, the ability of edge-adapted bats to use the edges may further be impaired by the lack of landscape complementation, i.e., the ability of different habitat types to provide the non-substitutable resources needed by bats, for instance roosting and foraging grounds (Dunning et al. 1999; Ethier & Fahrig 2011).

Although the effects of edge density negatively affected bat diversity at different levels, the effects of the number of patches were positive for total bat activity. In the Kenyir Lake, the islands were relatively well connected, interpatch distance rarely exceeding 1 km: considering that the home range of many bats making up the local assemblage exceeds this distance (Wilson et al. 2016), it is therefore likely that most bats used multiple islands as stepping stones to commute over the lake, despite the high inhospitality of the water matrix (Albrecht et al. 2007). It is also possible that a higher number of patches in the landscape was associated with a higher overall structural complexity, a landscape characteristic known to favour the activity of edge foragers (Ewert et al. 2023), or with a higher landscape complementation, each island potentially offering complementary resources (Dunning et al. 1999; Fahrig 2019). Indeed, in Kenyir Lake, the most deforested landscapes are mainly covered with water, and unlike patch expansion, any patch addition contributes to enhancing the landscape’s heterogeneity, thereby creating a “patchwork effect”. By allowing the presence of disturbance-adapted species, this patchwork effect characterised by a reduced inter-patch distance as well as an increased functional connectivity and habitat complementation, might therefore promote bat activity (Palmeirim et al. 2019).

### 4.2 Direct effects of habitat amount

Our results emphasise the existence of a positive direct effect of forest cover on sonotype richness. Positive effects of habitat amount on richness are widespread and observed on a variety of taxa when considered at the landscape-scale, including arthropods (With & Payne 2021), birds (Torrenta & Villard 2017), and small mammals (Merckx et al. 2019; Palmeirim et al. 2019). Other studies on bats in tropical landscapes have reported similar findings in both terrestrial (Muylaert et al. 2016) and insular fragmented landscapes (Meyer & Kalko 2008). Yet, we stress that we chose to use sonotype richness in our study: unlike species richness, there can be a lot of variability in the number of species represented by each sonotype: here, sonotype richness should therefore be interpreted more as a measure of functional diversity (i.e., how many different foraging strategies are represented) rather than a true reflection of species diversity.

In line with our hypotheses, the direct effects of habitat amount were positive on forest sonotype activity. FM and CF calls used by forest bats are suited to high levels of vegetation clutter where other sonotypes are not able to distinguish prey from the background (Denzinger & Schnitzler 2013; Schnitzler & Kalko 2001). Accordingly, Núñez et al. (2019) identified CF species as being particularly dependent on forest cover, with CF bats being less abundant in clearings and edges compared to forest interiors. Furthermore, most species classified as forest species based on their echolocation design also had a typical forest-adapted morphology comprising wide and short wings (low AR and WL) (Senawi & Kingston 2019). Allowing an agile but slow flight, these characteristics allow bats to navigate in highly cluttered forested areas but make long distance flights energetically costly and increases their risk of predation when flying over open spaces (Altringham 2011; Bader et al. 2015). In that sense, the uniform inhospitality of the water matrix likely plays a great role in the response of assemblage composition: the ability of a species to exploit different types of matrix habitat largely relies on species ecomorphological traits (Bader et al. 2015; Farneda et al. 2015). Through the reduction of forest commuting area, forest species may be unable to forage in high contrast matrices such as open water (Meyer & Kalko 2008) or agricultural land (Rocha et al. 2017).

### 4.3 Habitat loss and fragmentation: two interrelated processes

There is compelling empirical evidence from patch-scale studies regarding the strong negative effects of fragmentation on biological communities (Chase et al. 2020; Haddad et al. 2015). This trend also holds true for bats (López-Bosch et al. 2021), including those at Kenyir Lake, where the activity of forest specialists was reduced in small, isolated islands (Hazard et al. 2023). Some have questioned the legitimacy of such local-scale approaches (e.g. Fahrig (2013, 2017), but see Fletcher et al. (2023)) because of their inability to disentangle patterns arising from habitat loss versus those attributed to habitat fragmentation *per se*. Despite the effort of landscape scale studies aimed at unravelling the respective effects of these forces, most have overlooked the inherent reality that in natural environments, nearly every alteration in habitat amount influences habitat configuration (Figure 1). Indeed, our results underscored habitat amount as being a strong predictor of habitat configuration: landscapes with greater forest cover consistently exhibited higher edge density and fewer patches in comparison to those with lower forest cover (Clément et al. 2017; Püttker et al. 2020; Villard & Metzger 2014). As such, given the influence of habitat configuration on the bat assemblage, we observed strong indirect effects of habitat amount mediated through configuration.

The indirect effects of forest cover primarily operated through changes in edge density rather than the number of patches. Indeed, although both configuration variables were influenced by forest cover, the impact of *edge* on overall bat diversity was more evident than that of number of patches, presumably because of the increased habitat heterogeneity favoured by a high number of patches, which would benefit edge and open-space foragers (Fahrig 2019). Accordingly, using multi-taxa pathway analysis, Püttker et al. (2020) accounted for the indirect effects of habitat loss through fragmentation over the whole gradient of habitat amount. Yet, the indirect effects they found were comparatively weaker than the ones we observed. In our case, the emergence of indirect effects of habitat amount resulted from 1) the strong effects of edge density on the response variables, and 2) the relationship between habitat amount and configuration being notably strong at low habitat amounts (Villard & Metzger 2014). Indeed, on average our landscape harboured less than 30% of forest cover, a portion of the habitat gradient where each increase in habitat amount is tightly associated with an increase in edge density, thereby explaining the strong relationship linking these variables (Figure 4) (Liu et al. 2016; Pickell et al. 2016).

Although direct effects of forest cover on bat sonotype richness were positive, indirect effects of forest cover on richness, total activity, and the activity of edge foragers were negative when mediated through edge density. These negative indirect effects also find their source in the non-linear interdependence between habitat amount and configuration. Indeed, the relationship linking *cover* and *edge* was bell shaped (Table 1, Figure S4): under 30% of forest cover, increased habitat amount caused a strong increase in edge density, a trend that did not hold in landscapes harbouring a higher amount of forested area. Consequently, given the negative influence of edge density on sonotype richness, total activity, and the activity of edge foragers (Figure 4, S3), indirect effects of habitat amount mediated through edge density were negative. Nevertheless, our results should be interpreted with caution as our sampling sites mostly covered highly deforested landscapes. Indeed, across the selected scales, landscapes were consistently highly deforested (Table 1). At intermediate and high habitat amounts, where the relationship between habitat amount and edge density is negative, the total effect of habitat amount mediated through configuration becomes positive (Villard & Metzger 2014). When separating landscapes between low (>30%), intermediate (30-60%) and high (>60%) habitat amounts to account for this non-linear relationship, Püttker et al. (2020) reported positive indirect effects of habitat loss outweighing their direct effects. They highlighted that in highly forested landscapes, the positive effects of habitat amount were not direct but were mainly caused by the associated decrease in edge density. When the habitat left in the landscape is relatively low (< 30%), minimising further habitat loss should remain a top conservation priority to preserve positive direct effects of habitat amount by preventing the occurrence of unwanted, small habitat amount-mediated edge effects. Conversely and along with patterns observed by Püttker et al. (2020), we observed that although weaker than their effects on *edge*, effects of forest cover on *n.patches* were negative. Resulting from the positive influence of *n.patches* on total activity, the indirect effects of *cover* mediated through *n.patches* were therefore negative on total activity. This result may stem from the fact that in highly deforested landscapes, any patch may have served as a stepping stone for bats to commute over the lake. If well connected, a myriad of small patches may therefore have been more accessible than few larger but isolated patches. In fact, some animals may paradoxically move more frequently between patches in degraded habitats, potentially explaining the negative indirect effects of habitat amount as mediated by number of patches on bat activity (Bélisle 2005; de la Peña-Domene & Minor 2014).

### 4.4 Conservation implications

Given the interdependence of habitat loss and fragmentation in real landscapes, these two processes cannot be managed independently, as previously thought (Smith et al. 2011). In highly deforested landscapes, restoring the forest cover would primarily involve the creation of edges rather than core habitat. Yet, in high-contrast matrixes, such edges can only support an impoverished diversity. We therefore suggest that conservation efforts should be concentrated towards minimising habitat loss above the context-specific threshold where increases in habitat amount become positively associated with edge creation. To maximise bat diversity in highly contrasted landscapes with overall low forest cover, conservation actions should promote the increase of the habitat amount in the landscape while (1) minimising any increase in the edge density, and (2) also promoting the increase of the number of patches. Given the positive effects of forest cover and the number of patches, habitat restoration should primarily consist of the creation of additional forest patches (Saura et al. 2014), rather than thin and long linear vegetation strips. Additionally, strategies could be based on increasing the size of existing patches in a way that edge length is minimised. Moreover, further deforestation should primarily be avoided in landscapes already characterised by low habitat amounts. In flat tropical lowlands, the construction of dams goes hand in hand with the creation of a myriad of small islands, thereby favouring the creation of edge-dominated deforested landscapes. We stress that flat areas must be avoided at all costs for dam implementation, to minimise the creation of landscapes that are so altered that even disturbance-adapted species avoid them. If taken together, these considerations would help to effectively maximise bat diversity across lowland tropical forests.

## 5 Conclusion

The ongoing debate surrounding the impacts of habitat loss versus habitat fragmentation has been a focal point of discussion within the theoretical realms of ecology and conservation. However, an essential, yet overlooked factor in this debate, is the intricate link between habitat amount and configuration. Here, we acknowledged the association between these forces to highlight that in heavily deforested landscapes, the strength and direction of the effects of habitat amount were heavily dependent on habitat configuration. This complex relationship underscores the necessity for an integrated conservation approach that considers how actions taken to address habitat loss can significantly influence edge creation, as well as overall habitat connectivity. Given the pervasiveness of habitat loss and edge effects on bat communities, including on disturbance-adapted species, we advocate that efforts should be made to restore and preserve core forest habitats while minimizing the creation of new edges within these habitats.

## Acknowledgements

We extend our sincere gratitude to David López-Bosch and Ahmad Faizul Bin Zulkifli for their invaluable assistance during the data collection process. We also wish to express our appreciation to the Economic Planning Unit, Department of Prime Minister, Malaysia, for granting us permission to conduct our research, as well as to the Department of Wildlife and National Parks Peninsular Malaysia for permitting our work in Kenyir (JPHL&TN (IP):100-34/1.24 Jld 14(57)). Furthermore, we acknowledge the support received from various individuals and organizations that enabled this research.

AFP was supported by the Outstanding Postdoctoral Fellowship of the Southern University of Science and Technology (SUSTech) and by the European Union’s Horizon 2020 research and innovation program under grant agreement No. 854248. LG received support from the China Thousand Young Talents Program (K18291101) and was recognized as a Guangdong Government distinguished expert (K20293101), along with support from the Shenzhen Government (Y01296116). JSPF’s research was funded by the Leverhulme Trust through an early-career fellowship (Award Reference: ECF-2020-571). NY’s contributions were supported by the UK’s Natural Environmental Research Council (NERC) through an EnvEast DTP scholarship (NE/L002582/1). Lastly, JS received support from the National Science Fund - USA through Texas Tech University (ST-2019-006) and the Malaysia Ministry of Higher Education (FRGS/1/2020/WAB11/ UKM/02/3).

