## Supplemental Material for "Indirect effects of habitat amount mediated by habitat configuration determine bat diversity at the landscape-scale in Peninsular Malaysia"

**Supplementary material**

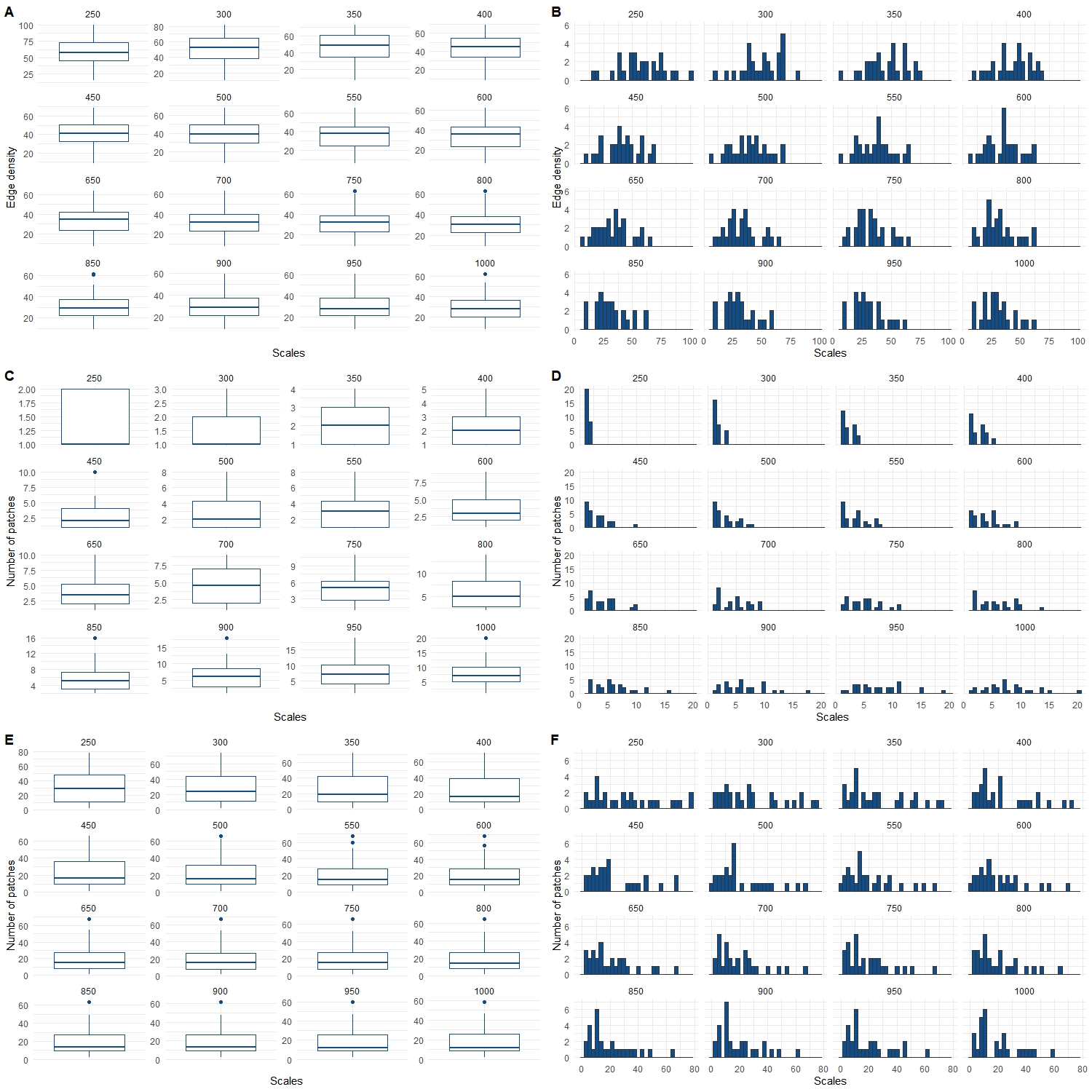

**Fig S1** Distribution of the variable “edge density” (A-B), “number of patches” (C-D), and “ percentage of forest cover” (E-F) over the different circular landscape scales (250 – 1000 m radii).

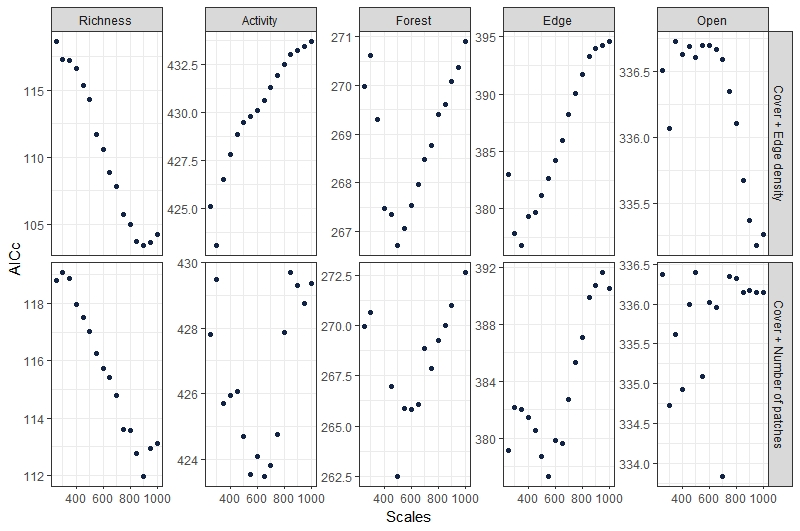

**Fig S2** determination of the scale of effect: for each model, the scale of effect was the one giving the lowest AICc when fitting a GLM response ~ cover + ed (first line), and response ~ cover + np (2nd line)

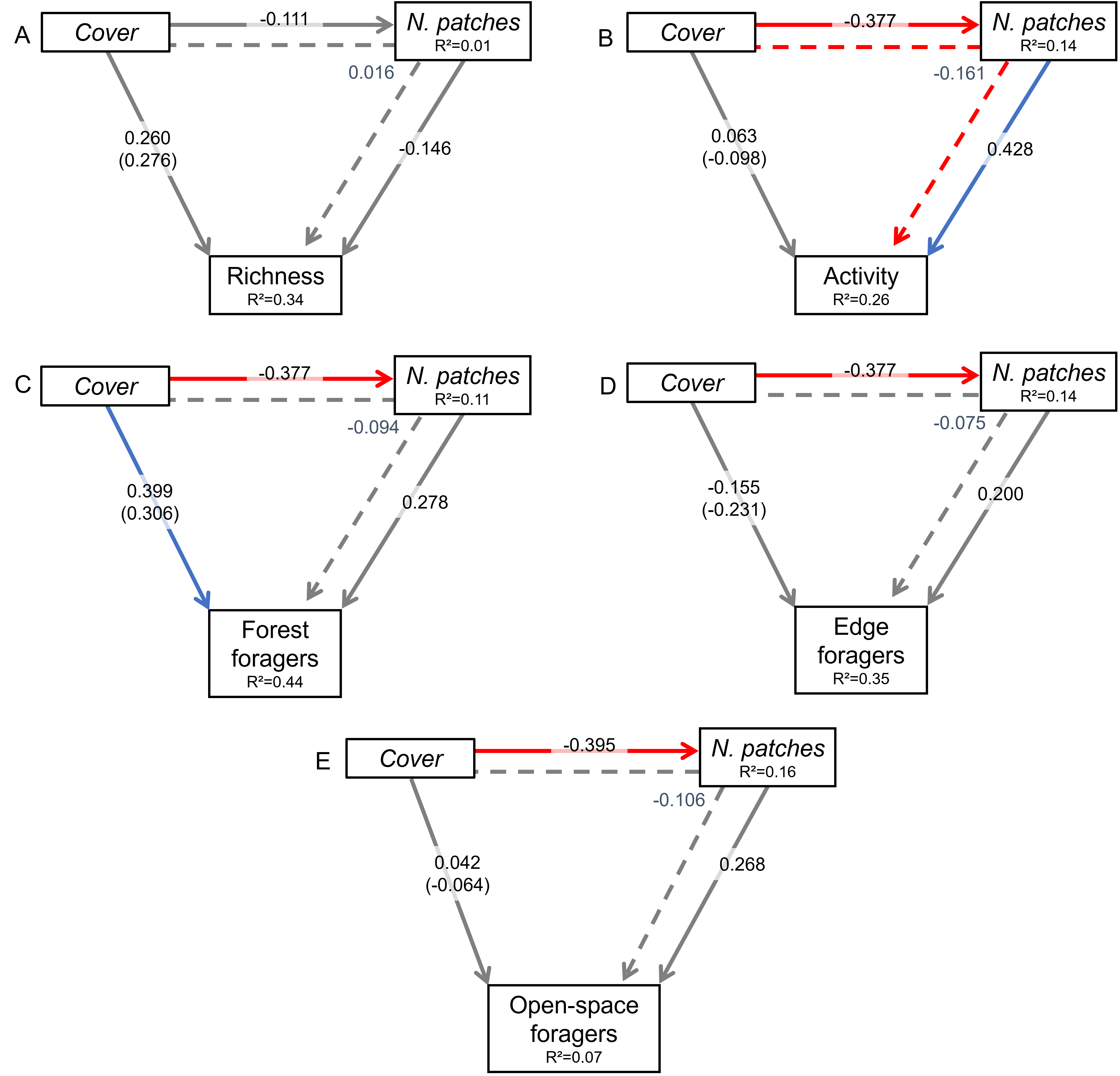

**Fig S3:** Results of the Piecewise Structural Equation Models representing the effects of habitat amount (*cover*) and *n.patches* on (a) sonotype richness, (b) total activity (log_10_ *x*), and activity of (c) forest sonotypes (log_10_ *x*), (d) edge sonotypes (log_10_ *x*), and (e) open-space sonotypes (log_10_ *x*). Blue arrows depict positive relationships, red arrows depict negative relationships, and grey arrows depict non-significant relationships. Solid arrows depict direct relationships, and dashed arrows represent indirect relationships. For each response variable, we indicated the standardised path coefficients. The distance point-edge was omitted from clarity as it was never significant. Coefficients for this variable can be found in Table S3.

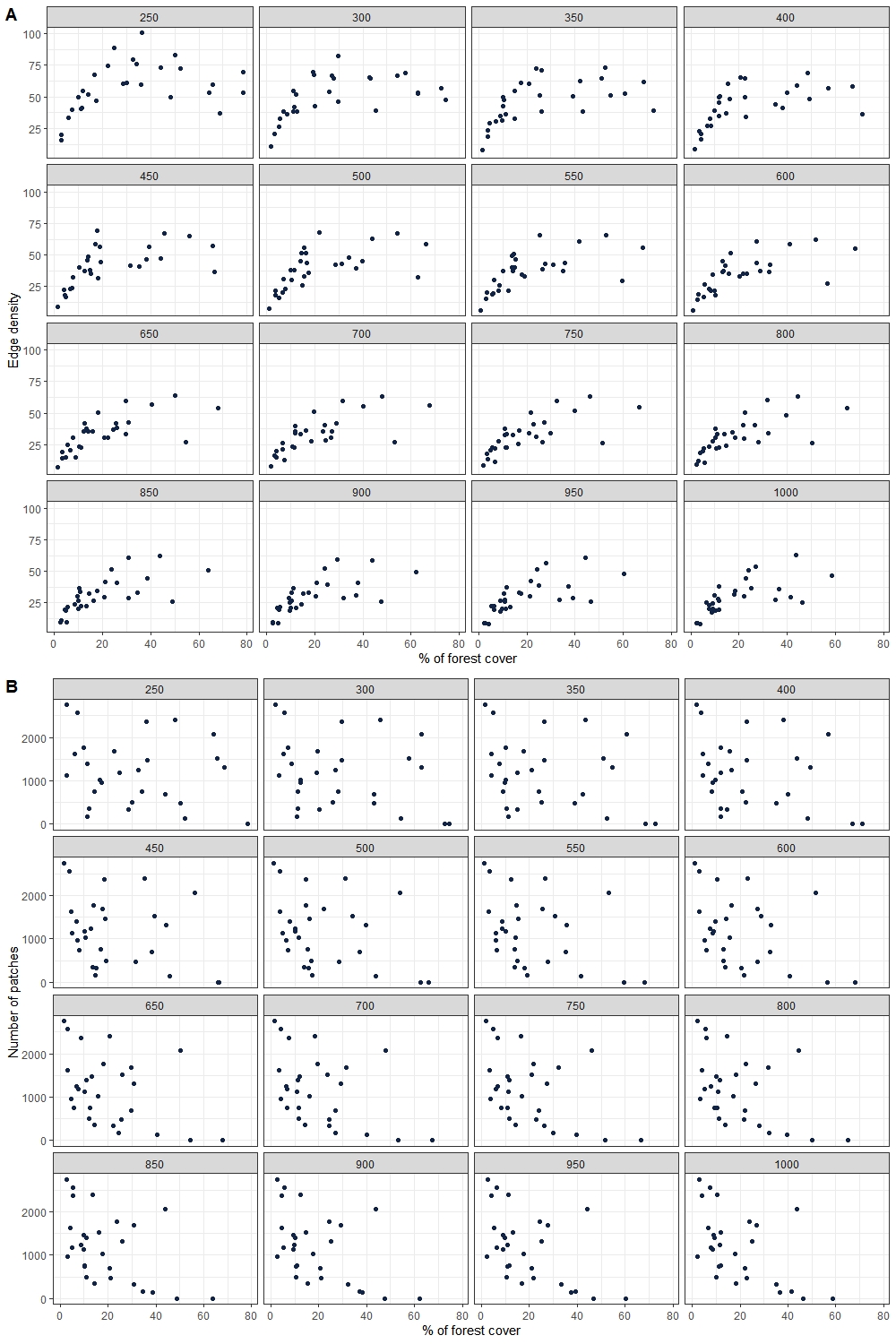

**Fig S4** Relationship between A) amount of forest cover and edge density and B) amount of forest cover and number of patches for 250 - 1000 m radii circular landscapes.

**Table S1.** Description of the acoustic activity recorded across the Kenyir Lake. For each guild, we provide details on the associated sonotypes and total activity (i.e., number of bat passes). Any bat passes that remained unidentified during the surveys were categorized as "unknown”. For further information on the potential species making up the sonotype, refer to Hazard et al. (2023).

| **Guild** | **Sonotype** | **Number of bat passes** |
| --- | --- | --- |
| Forest | *H. diadema* | 392 |
|  | *R. affinis* | 146 |
|  | *R. trifoliatus* | 2268 |
|  | *R. luctus* | 6 |
|  | *R. refulgens* | 2 |
|  | *H. kunzi* | 1 |
|  | *H. cervinus* | 1 |
|  | *H. bicolor* | 2 |
|  | CF.46 | 1 |
|  | FM | 35 |
| Open-space | LF | 2048 |
|  | FMqCF2 | 162 |
|  | FMqCF3 | 1441 |
| Edge | FMqCF4 | 13 195 |
|  | FMqCF5 | 389 |
|  | QCF | 119 |
| Unknown | Unknown | 988 |

**Table S2:** Piecewise SEM’s path coefficients and the associated p values of the relationship between each response variable and the distance between the detector and forest edge (ED).

|  |  | | **Model** |
| --- | --- | --- | --- |
| Effect | Response | Predictor | Richness ~ Cover + *Edge* + *Near.dist* |
| Direct | Richness | ED | -0.422 ± 0.151  (-0.690 - -0.102) |
|  | Richness | %cover | 0.476 ± 0.147  (0.197 - 0.736) |
|  | Richness | Dist.edge | -0.143 ± 0.130  (-0.135 - 0.386) |
|  | ED | %cover | 0.516 ± 0.118  (0.265 - 0.742) |
|  | ED | %cover² | -0.332 ± 0.129  (-0.632 - -0.093) |
| Indirect | Richness | %cover | -0.218 ± 0.102  (-0.425 - -0.425) |
|  | Richness | %cover² | 0.140 ± 0.083  (0.018 - 0.018) |
|  |  |  | **Activity ~ Cover + ED** |
| Direct | Activity | ED | -0.553 ± 0.126  (-0.741 - -0.263) |
|  | Activity | %cover | 0.062 ± 0.138  (-0.239 - 0.306) |
|  | Activity | Dist.edge | -0.025 ± 0.149  (-0.304 - 0.279) |
|  | ED | %cover | 0.710 ± 0.094  (0.461 - 0.839) |
|  | ED | %cover² | -0.605 ± 0.098  (-0.765 - -0.385) |
| Indirect | Activity | %cover | -0.392 ± 0.108  (-0.587 - -0.182) |
|  | Activity | %cover² | 0.334 ± 0.100  (0.150 - 0.534) |
|  |  |  | **Forest ~ Cover + ED** |
| Direct | Forest | ED | 0.163 ± 0.193  (-0.222 - 0.509) |
|  | Forest | %cover | 0.221 ± 0.162  (-0.104 - 0.526) |
|  | Forest | Dist.edge | 0.098 ± 0.160  (-0.229 - 0.406) |
|  | ED | %cover | 0.592 ± 0.114  (0.383 - 0.813) |
|  | ED | %cover² | -0.445 ± 0.140  (-0.740 - -0.208) |
| Indirect | Forest | %cover | 0.097 ± 0.113  (-0.107 - 0.332) |
|  | Forest | %cover² | -0.073 ± 0.088  (-0.293 - 0.065) |
|  |  |  | **Edge ~ Cover + ED** |
| Direct | Edge | ED | -0.520 ± 0.124  (-0.740 - -0.288) |
|  | Edge | %cover | -0.123 ± 0.118  (-0.387 - 0.079) |
|  | Edge | Dist.edge | -0.053 ± 0.102  (-0.229 - 0.177) |
|  | ED | %cover | 0.661 ±0.112  (0.420 - 0.839) |
|  | ED | %cover² | -0.545 ± 0.125  (-0.768 - -0.298) |
| Indirect | Edge | %cover | -0.343 ± 0.119  (-0.590 - -0.182) |
|  | Edge | %cover² | 0.283 ± 0.101  (0.130 - 0.555) |
|  |  |  | **Open ~ Cover + ED** |
| Direct | Open | ED | -0.211 ± 0.198  (-0.557 - 0.217) |
|  | Open | %cover | 0.022 ± 0.209  (-0.345 - 0.476) |
|  | Open | Dist.edge | 0.045 ± 0.190  (-0.355 - 0.387) |
|  | ED | %cover | 0.664 ± 0.109  (0.446 - 0.842) |
|  | ED | %cover² | -0.540 ± 0.09  (-0.769 - -0.305) |
| Indirect | Open | %cover | -0.140 ± 0.132  (-0.416 - 0.097) |
|  | Open | %cover² | 0.114 ± 0.113  (-0.064 - 0.379) |

**Table S3:** Piecewise SEM’s path coefficients and the associated p values of the relationship between each response variable and the distance between the detector and forest edge (ED). ED: Edge density, NP: number of patches.

|  |  | | **Model** |
| --- | --- | --- | --- |
| Effect | Response | Predictor | **Richness ~ Cover + ED** |
| Direct | Richness | NP | –0.146 ± 0.147  (–0.437 - 0.142) |
|  | Richness | %cover | 0.260 ± 0.186  (–0.109 - 0.595) |
|  | Richness | Dist.edge | 0.224 ± 0.150  (–0.067 - 0.515) |
|  | NP | %cover | –0.111 ± 0.149  (–0.385 - 0.205) |
| Indirect | Richness | %cover | 0.016 ± 0.188  (0.033 - 0.018) |
| Direct | Activity | NP | 0.428 ± 0.142  (0.142 - 0.690) |
|  | Activity | %cover | 0.063 ± 0.171  (–0.278 - 0.390) |
|  | Activity | Dist.edge | –0.135 ± 0.172  (–0.452 - 0.214) |
|  | NP | %cover | –0.377 ± 0.127  (–0.596 - –0.093) |
| Indirect | Activity | %cover | -0.161 ± 0.078  (-0.349 - -0.33) |
| Direct | Forest | NP | 0.278 ± 0.191  (–0.122 - 0.607) |
|  | Forest | %cover | 0.399 ± 0.150  (0.046 - 0.642) |
|  | Forest | Dist.edge | 0.086 ± 0.140  (–0.193 - 0.349) |
|  | NP | %cover | –0.337 ± 0.114  (–0.542 - –0.095) |
| Indirect | Forest | %cover | –0.094 ± 0.073  (–0.261 - 0.025) |
| Direct | Edge | NP | 0.200 ± 0.191  (–0.244 - 0.514) |
|  | Edge | %cover | –0.155 ± 0.164  (–0.437 - 0.223) |
|  | Edge | Dist.edge | –0.198 ± 0.189  (–0.572 - –0.572) |
|  | NP | %cover | 0.377 ± 0.127  (–0.596 - –0.095) |
| Indirect | Edge | %cover | –0.075 ± 0.078  (–0.231 - 0.071) |
| Direct | Open | NP | 0.268 ± 0.177  (–0.107 - 0.576) |
|  | Open | %cover | 0.042 ± 0.222  (–0.381 - 0.472) |
|  | Open | Dist.edge | –0.003 ± 0.177  (–0.323 - 0.367) |
|  | NP | %cover | –0.395 ± 0.111  (–0.587 - –0.155) |
| Indirect | Open | %cover | –0.106 ± 0.076  (–0.272 - 0.031) |
